# Inhibition of β-catenin signaling by Amyloid-β in endothelial cells impairs vascular barrier integrity

**DOI:** 10.1101/2024.11.05.622172

**Authors:** Joseph W. Leasure, Thi Mai, Avi Fogel, Pierina Danos, Li Wang, Mark A. Sanborn, Zijing Ye, Young-Mee Kim, Xinhe Wang, Jennifer Flores, Muzamil Mushtaq, Peter T. Toth, Sang Ging Ong, Orly Lazarov, Jalees Rehman

**Affiliations:** Department of Biochemistry and Molecular Genetics, University of Illinois, College of Medicine, Chicago, IL 60607; Research Resources Center, University of Illinois at Chicago, Chicago, IL 60612; Department of Pharmacology and Regenerative Medicine, University of Illinois, College of Medicine, Chicago, IL 60612; Department of Anatomy and Cell Biology, University of Illinois, College of Medicine, Chicago, IL 60612; Department of Biomedical Engineering, University of Illinois at Chicago, Chicago, IL 60612; Division of Cardiology, Department of Medicine, University of Illinois, College of Medicine, Chicago, IL 60612

**Keywords:** endothelium, endothelial cells, Alzheimer’s disease, Amyloid-beta, induced pluripotent stem cells, β-catenin, blood-brain barrier

## Abstract

Dysfunction of the blood-brain barrier (BBB) is emerging as a critical mediator of Alzheimer’s Disease (AD) progression and precedes the formation of large Amyloid-β (Aβ) plaques in AD patients. Maintenance of the blood brain barrier integrity by brain microvascular endothelial cells (BMECs) is essential for brain homeostasis and is in part maintained by WNT/β-catenin signaling. We hypothesized that Aβ induces blood-brain barrier dysfunction by inhibiting WNT/β- catenin signaling in brain endothelial cells, leading to reduced expression of BBB-related genes and barrier function. In brain endothelial cells, we found that Aβ directly decreased β-catenin transcriptional activity, but also inhibited activation of β-catenin by the WNT3A peptide. Additionally, Aβ reduced barrier integrity of ECs, which was in part restored by exposure to exogenous WNT3A. We observed Aβ-induced inhibition within 3 hours of exposure. Aβ-induced inhibition of β-catenin signaling was consistent across both immortalized brain ECs and ECs differentiated from human induced pluripotent stem cells(iPSCs). We next investigated human iPSCs with and without the N141I mutation in presenilin-2 (*PSEN2*), known to increase Aβ buildup and familial AD risk. We found the AD ECs showed smaller cell size and expressed lower VE-cadherin at the protein level. RNA sequencing revealed that the presence of *PSEN2* N141I did not significantly impact gene expression of genes coding for proteins canonically involved in barrier integrity but did show differences in the expression of transposable elements. These results indicate that BBB dysfunction early in AD may be the result of Aβ in the brain impairing WNT/β-catenin signaling which maintains the barrier integrity of brain endothelial cells.

## INTRODUCTION

Several studies suggest that blood-brain barrier (BBB) dysfunction promotes the development of neurodegenerative diseases^1–4^. BBB dysfunction is characterized by increased permeability of the brain vascular endothelium,^1, 2, 5^ and aging is strongly associated with increased BBB permeability^2, 5–8^. In homeostatic conditions, the brain vascular endothelium maintains a high level of barrier integrity to prevent the entry of toxins, pathogens or excessive immune cells into the brain parenchyma^2, 7, 9^. Conversely, increased permeability in the setting of disease allows for the entry of immune cells, reduced efflux of toxic peptides and molecules, and imbalance of solute and ion transport that reduce neuronal health^2, 9, 10^

Disruption of the blood-brain barrier is also emerging as a pathogenic mechanism in the development of Alzheimer’s Disease (AD)^4, 11, 12^. Investigation into AD has focused primarily on the neurotoxic peptides Amyloid-Beta (Aβ) and hyperphosphorylated tau, two pathological hallmarks of AD thought to be causally involved in the disease progression^13, 14^. Neurons produce amyloid precursor protein (APP), a small protein that is ultimately processed into neurotoxic Aβ by γ-secretase^14, 15^. Secreted Aβ accumulates to form oligomers and large plaques in the brain that which promote cognitive decline and neurodegeneration in AD^14^. Heritable mutations in the genes that code for the proteins that make up the γ-secretase complex, such as presenilin-1 (*PSEN1*) and presenilin-2 (*PSEN2*), are associated with increased Aβ generation and earlier onset of heritable AD^16^. Notably, some mutations, such as the N141I mutation in *PSEN2*, do not increase overall Aβ generation, but instead increase the ratio of neurotoxic Aβ_1-42_ to non-toxic Aβ_1-40_^17^. However, it is not clear whether these peptides contribute to AD progression via the increased dysfunction of the BBB^1, 2, 9^

Several mechanisms may account for how Aβ may disrupt the BBB. The first is a notable correlation between Aβ and reduced WNT/β-catenin signalling^12, 18–21^. Disrupted β-catenin signaling, seen through decreased expression of WNT ligands and increased markers of β- catenin suppression such as GSK3β, have been proposed as important mediators of AD development ^19, 20, 22^. A number of cellular mechanisms describing the direct effect of Aβ have been described, including the binding of Aβ to neuronal Frizzled receptors which prevents the binding of WNT ligands, thus reducing neuroprotective WNT/β-catenin signaling^18, 19^ but there is little known about the effects of Aβ on endothelial β-catenin signaling. This provides an intriguing connection to vascular dysfunction in AD, as brain endothelial cells rely on high WNT/β-catenin signaling to maintain their unique BBB phenotype^23–26^, and restoring such signaling in mouse and cell models have been shown to restore endothelial barrier function^12^.

In this study, we utilize both the human CMEC/D3 brain endothelial cell line and iPSC-derived endothelial cells to investigate the effect of Aβ exposure on endothelial β-catenin signaling and the mediators of endothelial barrier integrity. We found Aβ acutely inhibited both endogenous β- catenin and WNT-induced upregulation of β-catenin targets. We also found Aβ reduced the expression of endothelial VE-cadherin, as well as reducing barrier integrity.

## RESULTS

### Amyloid-β inhibits endogenous β-catenin activity and the WNT3A-induced upregulation of β- catenin target genes

We first examined the effects of exogenous Amyloid-β exposure on hCMEC/D3 brain endothelial cells at two concentrations in the media: 0.5 µM and 1 µM for 24 hours. The negative controls were instead given 24 hours of 1 μM scrambled Aβ peptide (“SCR”). The 24 hour exposure time was chosen to emulate a long-term exposure to Aβ, as we would expect in a patient over the course of AD development. The isolated RNA was converted to cDNA and used in qPCR to determine changes to β-catenin transcription targets *AXIN2* and *TCF7*. Beta- 2-Microtubilin was also measured as a housekeeping gene control. **(Fig 1A)** Comparing the SCR control expression to Aβ exposed samples, exposure to Aβ results in decreased expression of both *AXIN2* and *TCF7* in a dose-dependent manner. Even 0.5 µM resulted in notable decreases in both genes, though the difference was not statistically significant for *AXIN2*, indicating the inhibitory effect was stronger against *TCF7* expression. However, at 1 µM both *AXIN2* and *TCF7* were reduced to almost 60% of its original expression compared to the SCR control. First, this data demonstrates the inhibitory and dose-dependent inhibition of Aβ. Secondly, for experiments going forward, this data indicates 1 µM as a useful experimental concentration where the effect on β-catenin is significant, as higher concentrations of Aβ could affect other pathways or cell health and confound the results^27^.

**Figure 1.**
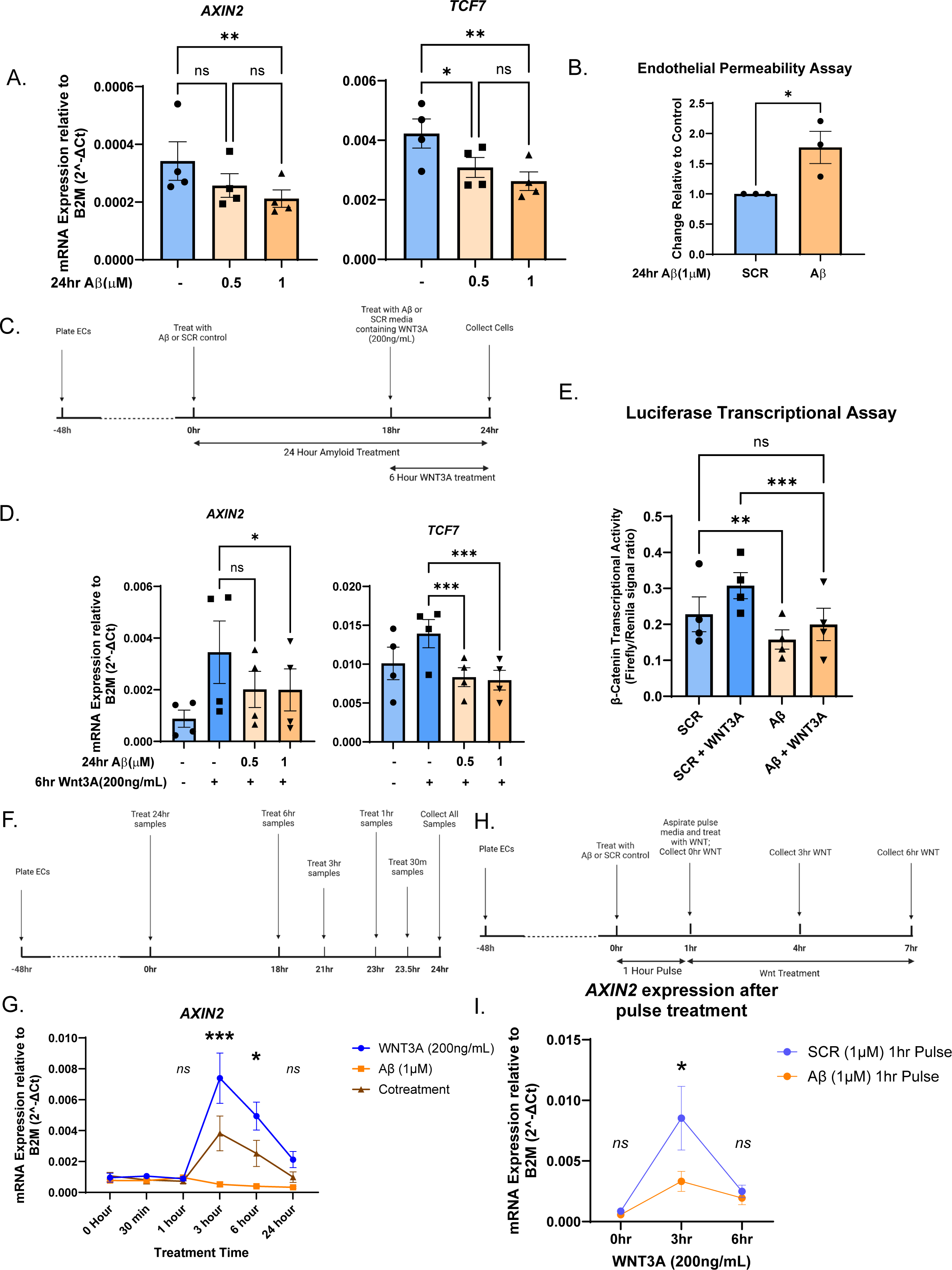
Amyloid-β and WNT3A exposure in *in vitro* hCMEC/D3 brain endothelial cells. A) RT-qPCR against β-catenin targets *AXIN2* and *TCF7* from brain endothelial cells exposed to 0.5µM and 1µM of Amyloid-Beta. Untreated cells were exposed to 1µM scrambled peptide control (SCR). Data is represented as a mean +/-SEM of n=4 independent experiments. Statistics were analyzed by one-way ANOVA followed by post-hoc Bonferroni’s multiple comparisons test. *AXIN2*: 0μM vs. 0.5μM *ns*=0.0653, 0.5μM vs. 1μM *ns p*=0.4499, *\*\*p=*0.0097. *TCF7: *p*=0.0365, *\*\*p=*0.0075, *ns p=*0.6072. B) Endothelial permeability assay recording the barrier permeability of brain endothelial cells exposed to either SCR or Aβ in the absence or presence of WNT3A activation. Data is represented as the mean +/-SEM of n=3 independent experiment. Statistics were analyzed with student’s unpaired, two-tailed *t-*test, *\*p*=0.0446. C) Timeline of Aβ and WNT3A exposure used for experiments D and E. Created with Biorender.com D) RT-qPCR against β-catenin targets *AXIN2* and *TCF7* in brain endothelial cells activated by WNT3A peptides either in the presence of SCR or increasing concentrations (0.5μM or 1μM) of Aβ. Statistics were analyzed by one-way ANOVA followed by post-hoc Bonferroni’s multiple comparisons test. *AXIN2: *p*=0.0477, *ns p=*0.0502. *TCF7:* 0.5μM Aβ *\*\*\*p=*0.0002, 1μM Aβ *\*\*\*p=*0.0001 E) Luciferase gene expression assay data showing β- catenin-dependent luciferase expression in brain endothelial cells exposed to either SCR or Aβ in the absence or presence of WNT3A activation. Statistics were analyzed by one-way ANOVA followed by post-hoc Bonferroni’s multiple comparisons test. *\*\*p=*0.0058*, ***p=*0.0003, *ns p=*0.3538. F)Timeline of Aβ and WNT3A in time-course experiment. Created with Biorender.com G) Time course experiment showing changes in *AXIN2* expression via RT-qPCR over 24 hours of exposure to either WNT3A, Aβ, or a combination treatment. Statistics were analyzed by two-way ANOVA followed by post-hoc Bonferroni’s multiple comparisons between each treatment at each time point. 1 hour: WNT3A vs. Aβ *ns p>*0.999, WNT3A vs Cotreatment *ns p>*0.999, Aβ vs. cotreatment *ns p>*0.999. 3 hour: WNT3A vs. Aβ \*\*\*\**p<0.0001,* WNT3A vs. Cotreatment *\*\*\*p=*0.0002, Aβ vs. cotreatment *\*\*\*p*=0.0006. 6 hour: WNT3A vs. Aβ \*\*\*\**p*<0.0001, WNT3A vs. Cotreatment \**p=*0.0156, Aβ vs. Cotreatment \**p=*0.0390. H) Timeline of SCR/Aβ 1 hour pulse before WNT3A activation. Created with Biorender.com. I) RT-qPCR of WNT3A-induced *AXIN2* expression after 1 hour pulse of either Aβ or SCR. Data is represented as a mean +/-SEM of n=4 independent experiments. Statistics were analyzed by two-way ANOVA followed by post-hoc Bonferroni’s multiple comparisons between SCR and Aβ pulse-treated samples. *ns p>0.9999, *p=0.0170*.

### Exposure to Aβ impairs barrier integrity of brain endothelial cells

In order to determine how this modulation of β-catenin transcriptional activity may affect the brain endothelial function, we performed a transwell FITC-dextran assay to measure the leakiness of the cell monolayer after these treatments. We expect an endothelial monolayer with reduced barrier function to demonstrate higher FITC-dextran signal in the receiver well compared to healthy control. As shown in **Fig 1B**, we found that the changes to monolayer permeability largely mirrored the changes to transcriptional activity seen in the previous experiment: Aβ significantly increased the permeability of the cell monolayer, indicating decreased cell barrier function and health. The FITC-dextran signal was shown to increase over 1.5-fold compared to the SCR-treated control, which *in vivo* could be expected to result in decreased Amyloid clearance and general disruption to the BBB.

### Exogenous Aβ inhibits WNT3A-induced activation of the β-catenin pathway

We next wanted to investigate how the presence of Aβ in the media would affect activation of the β-catenin pathway by WNT3A. The experimental layout in **Fig 1C** was utilized, where the cells were pre-treated with either Aβ or SCR to replicate a long-term exposure to amyloid peptides. The cells were then treated with 200 ng/mL WNT3A in media that also contained Aβ or SCR before being collected for RT-qPCR and collected after 6 hours of WNT3A activation **(Fig 1D)**. Compared to the cells treated only with WNT3A, cells co-treated with both Aβ and WNT3a saw a notable reduction in the expression of both AXIN2 and TCF7. In the case of AXIN2, the expression levels after cotreatment were still increased over 2-fold compared to the SCR negative control, but much lower than the nearly 4-fold increase of WNT3A alone. On the other hand, the cotreatment reduced TCF7 expression to even lower than the levels seen with the SCR negative control.

To directly investigate transcriptional activity in response to WNT3A and Aβ, a luciferase assay was performed on endothelial cells transfected with β-catenin-dependent luciferase plasmid **(Fig 1E)**. The cells were treated with either Aβ, WNT3A, or a combination with the same treatment timing as the previous experiment. This experiment demonstrated a similar Aβ-dependent decrease of both endogenous β-catenin signaling on its own compared to the negative SCR- treated control. Notably, in this experiment the cotreatment of Aβ with WNT3A entirely blunted WNT3A’s activation of β-catenin-dependent transcriptional activity, bringing it to approximately the same level as the SCR treated negative control.

### Aβ exhibits inhibitory effect against WNT3A-induced AXIN2 expression over 24 hours

With the previous data, we were interested in seeing whether the inhibitory effect would persist if the treatments were given at the same time, instead of pre-treating with Aβ. To do this, we performed a time course treatment of either WNT3A, Aβ, or the combination treatment over the course of 24 hours as shown in **Fig 1F**. In this experiment, we found that endogenous *AXIN2* levels were suppressed after just 3 hours of exposure time and continued to decline up to the final 24 hours of exposure **(Fig 1G)**. At 24 hours, the *AXIN2* expression level of Aβ was approximately 42% of the expression at 0 hours of treatment. Looking specifically at the effect of Aβ had on WNT3A-induced expression changes, we also found that the upregulation of *AXIN2* that begins at 3 hours of WNT3A exposure is inhibited when the cells are simultaneously treated with Aβ peptide. This difference can be seen by 3 hours, where we first see a strong increase in AXIN2, but this phenomenon persists for at least 6 hours. The effect is still present at 24 hours of exposure, though it is no longer statistically significant. This demonstrates the effect of Aβ on the β-catenin pathway acts early and is not dependent on extended pre-treatment.

### Pulse treatment with Aβ inhibits subsequent WNT3A activation

To further investigate the timeframe of how Aβ effects β-catenin activation by WNT3A in endothelial cells, we next performed a pulse treatment experiment using a 1-hour pulse of 1 μM SCR or Aβ, before replacing the media entirely with media containing only WNT3A. The timeline of the experiment is clarified in **Fig 1H** and the cells were collected for RT-qPCR against *AXIN2*. In this experiment, we found that the pulse treatment with Aβ ultimately inhibited the activation of *AXIN2* by WNT3A at 3 hours when compared to the SCR pulse-treated control (**Fig 1I**). This surprising result demonstrates the potency of Aβ’s inhibitory effect against WNT3A, able to act antagonistically after only 1 hour of exposure. However, this effect was no longer present by 6 hours of WNT3A treatment, also demonstrating the limits of Aβ’s inhibitory effects on β-catenin activation by WNT3A.

### Aβ exposure inhibits β-catenin target expression in iPSC-derived ECs

We next decided to investigate whether the inhibitory property of Aβ persists across other *in vitro* models of ECs, specifically iPSC-derived ECs. The 273 iPSC line was differentiated into endothelial cells using a previously published protocol^28^ and differentiation efficacy was shown to be approximately 95% CD31+ endothelial cells **(Supp Fig 1)**. To investigate the effect of Aβ on endogenous β-catenin activity, mature differentiated ECs were exposed to either scrambled peptide control or Aβ at 1µM and collected at (**Fig 2A**) 6hours and (**Fig 2B**) 24 hours for RT-qPCR analysis. In addition to *AXIN2* and *TCF7*, two β-catenin targets directly involved in the β- catenin pathway, we measured the expression of *CLDN5*, a gene regulated by β-catenin that codes for the tight junction protein claudin-5. Aβ significantly reduced the expression level of *AXIN2* at 6 hours compared to the SCR control, a reduction sustained through 24 hours of exposure as well. The inhibition of *TCF7* expression by Aβ at 6 hours is also significant, but a more subtle effect compared to *AXIN2*. By 24 hours, there is no longer a statistically significant difference but the inhibition of *TCF7* is still present compared to control. Looking at the barrier function gene *CLDN5*, we can see that there is no statistically significant difference in expression after 6 hours of exposure, but at 24 hours, Aβ-exposed endothelial cells expressed lower *CLDN5* compared to control. These results support our previous data that exposing cells to Aβ can reduce the expression of β-catenin targets *AXIN2* and *TCF7*, though interestingly some targets, such as *CLDN5* take longer to demonstrate inhibition by the peptides. This may be the result of additional regulation by other pathways that modulate *CLDN5* such as VEGF^29^.

**Figure 2.**
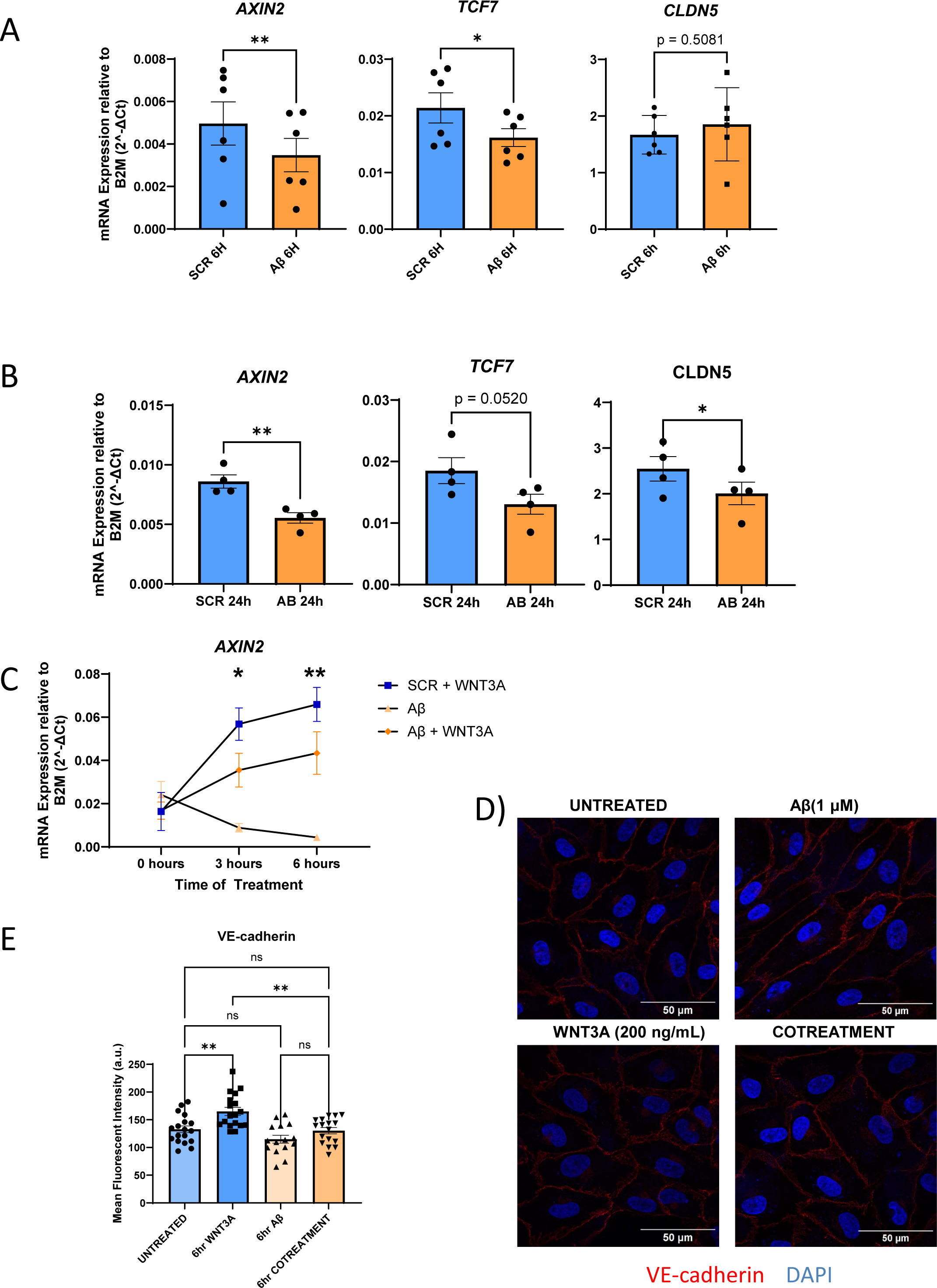
Amyloid-β and WNT3A exposure in *in vitro* iPSC-derived endothelial cells. A) RT-qPCR of endothelial cells exposed to either 1 μM SCR control or Aβ for 6 hours before collection. Data is represented as the mean +/-SEM of n=6 independent experiment. Statistics were analyzed with student’s paired, two-tailed *t-*test. *\*\*p=*0.0034, \**p=*0.0120. B) RT-qPCR of cells exposed to SCR control or Aβ for 24 hours. Data is represented as the mean +/-SEM of n=4 independent experiments. Statistics were analyzed with student’s paired *t-*test. *\*\*p=*0.0041, *\*p=*0.0126. C) Time course RT-qPCR data of *AXIN2* expression from iPSC-derived ECs exposed to either SCR (1 µM) and WNT3A (200ng/mL), Aβ (1 µM), or Aβ (1 µM) and WNT3A (200 ng/mL). Data is represented as a mean +/-SEM of n=4 independent experiments. Statistics were analyzed by two-way ANOVA followed by post-hoc Bonferroni’s multiple comparisons between each sample. 3 hours: SCR+WNT3A vs. Aβ *\*\*\*\*p<*0.0001, SCR+WNT3A vs. Aβ *ns p=*0.0310, Aβ vs Aβ+WNT3A *\*p=*0.0288. 6hours: SCR+WNT3A vs. Aβ *\*\*\*\*p<*0.0001, SCR+WNT3A vs. Aβ+WNT3A *ns p=*0.0779, Aβ vs Aβ+WNT3A *\*\*p=*0.0011. D) Representative immunofluorescent images of iPSC-ECs stained for VE-cadherin (red) expression after 6 hours of stimulation. DAPI is shown in blue. Scale bar: 50 μm. E) Quantified VE-cadherin fluorescent staining data of iPSC-ECs exposed to the listed treatments for 6 hours before fixation. Data is represented as a mean +/-SEM of independent experiments. UNTREATED and WNT3A n=18, Aβ n=15, COTREAMTENT n=17. Statistics were analyzed by one-way ANOVA followed by the post-hoc Bonferroni’s multiple comparisons shown. UNTREATED vs. WNT3A \*\**p=*0.0031, UNTREATED vs. Aβ *ns p=*0.3090, UNTREATED vs. COTREATMENT *ns p>*0.9999, WNT3A vs. COTREATMENT *\*\*p=*0.0015, Aβ vs. COTREATMENT *ns p=*0.5549.

### Aβ abrogates WNT3A-induced AXIN2 expression over 24 hours

We next decided to investigate whether Aβ inhibits WNT3A-induced upregulation of β-catenin targets, we decided to perform a time-course-RT-qPCR experiment to analyze changes to the expression of AXIN2 (**Fig 2C**). Consistent with previous data, exposure to Aβ decreased AXIN2 expression substantially, showing a distinct reduction at three hours of exposure that persisted after a total of six hours. Exposure to WNT3A, on the other hand, predictably increased expression of AXIN2 starting at three hours and persisted after six hours of exposure. When cotreated with both Aβ and WNT3A, the expression is notably reduced compared to the WNT3A-only samples at both three hours and six hours of exposure. This result is consistent with our previous experiments and demonstrates the short time frame across which Aβ can exhibit its inhibitory effects.

### WNT3A-induced increases of junctional VE-cadherin is inhibited by Aβ exposure

To determine how this inhibition may be affected by WNT3A stimulation, iPSC-ECs were exposed to Aβ, SCR+WNT3A, or Aβ+WNT3A cotreatment for 6 hours before being fixed and stained against VE-cadherin, the major component of endothelial adherens junctions. These were compared to untreated control cells. Representative images of the VE-cadherin signal found is shown in **Figure 2D** and is quantified in **Figure 2E**. We found that Aβ exposure resulted in a subtle decrease in the amount of VE-cadherin found at the junctions, which may partially explain the barrier integrity differences found in previous experiments. WNT3A exposure led to a substantial increase in the expression of VE-cadherin compared to SCR, which was significantly brought down when cotreating with Aβ to the level of untreated control. This data indicates that the presence of Aβ can counteract WNT3A-induced VE-cadherin expression. However, the more subtle differences between treatments compared to earlier barrier function experiments indicate that Aβ-induced dysfunction may involve more barrier genes that deserve to be investigated or may continue developing over time.

### Endothelial cells with PSEN2 N141I SNV/WT mutation show subtle changes to endothelial cell health in basal conditions

To investigate the effect of heritable familial Alzheimer’s Disease mutations on endothelial cell function, we generated endothelial cells from iPSCs carrying the fAD N141I mutation in the *PSEN2* gene. This single nucleotide variant (SNV) line was generated by CRISPR editing to possess one copy of the mutation (SNV/WT), and we also obtained the isogenic control REV/WT line which was generated by CRISPR-modifying the mutated allele back to wildtype (revertant or REV). Direct comparison of these two iPSC lines and the differentiated ECs allows for the investigation of the specific effects of the familial AD *PSEN2* mutation and the resultant increased generation of Aβ. Flow cytometry analysis shows that there was no difference in the efficiency of CD31+ ECs differentiated from either line, indicating that the cell lines can be utilized to compare their endothelial phenotype (**Supp Fig 1**). The PSEN2 N141I mutation is associated with increased deposition of Aβ_1-42_ relative to Aβ_1-40_^30^, thus we used ELISA against Aβ_1-42_ to quantify difference in Aβ generation in these iPSC-derived ECs (**Fig 3A**). We found that there was a trend towards increased Aβ generation with each differentiated pair showing more Aβ generation in the SNV/WT line.

**Figure 3.**
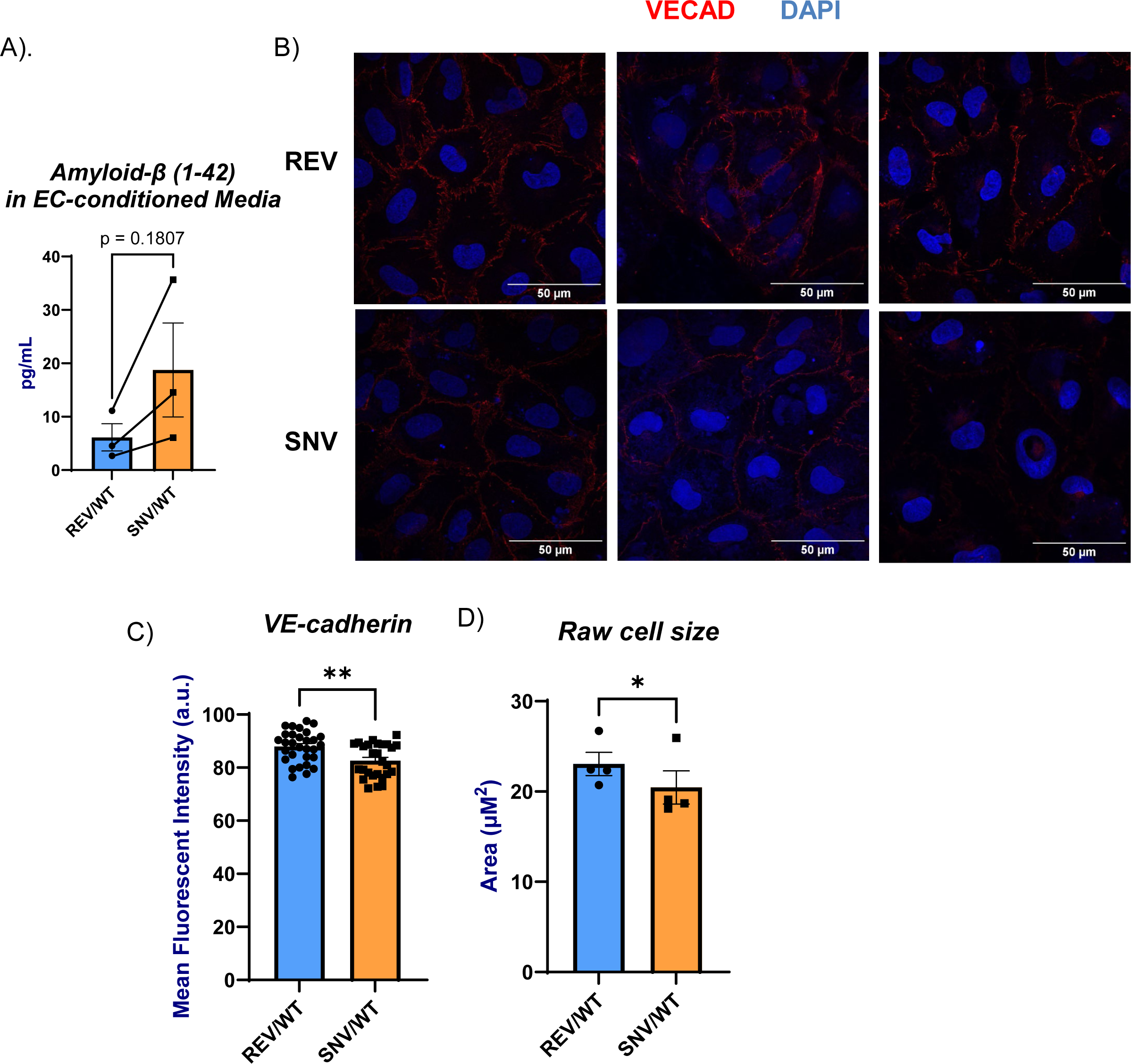
Comparison of iPSC-derived endothelial cells possessing the *PSEN2* N141I mutation (SNV/WT) compared to control cells with the mutation corrected (REV/WT). A) Expression of Aβ(1-42) in 72 hour conditioned media from either REV/WT or SNV/WT endothelial cells, as measured by ELISA. Lines were drawn to show the relationship between samples within each replicate. Statistics were analyzed with student’s paired, two-tailed *t-*test. B) Representative fluorescent images of confluent iPSC-derived ECs stained for VECAD (red) between REV/WT and SNV/WT cell lines. Scale bar = 50μM. Average quantification is shown in Fig 3D. C) Average cell size of confluent iPSC-derived REV/WT and SNV/WT endothelial cells, as measured from the fluorescent cell images described in figures 3C and 3D. Statistics were analyzed as n=4 independent experiments with student’s paired, two-tailed *t-*test. *\*p=0.0414.* D) Quantification of the fluorescent intensity of stained VECAD at cell barriers from the previous experiment. Data is represented as the mean of independent experiments. REV/WT n=30, SNV/WT n=27. Statistics were analyzed with student’s paired, two-tailed *t-*test.

We next plated the cells at equal cell concentrations onto gelatin-covered coverslips and grew both lines to confluence before fixing the cells at the same time for immunofluorescent analysis. We stained for VE-cadherin in order to visualize the size of the endothelial cells and adherens junctions. The fluorescent intensity of stained VE-cadherin at the cell borders was significantly weaker in the fAD SNV/WT cells compared to the REV/WT (**Figure 3B and 3C**). The VE-cadherin signal at the junctions was used to manually measure the average cell size of differentiated ECs. We found that the REV/WT control cells were larger on average, possibly a sign of better cell health (**Figure 3D**). This indicates that the paired control REV/WT and fAD SNV/WT cells can both be successfully differentiated into endothelial cells that can be compared *in vitro* to investigate the *PSEN2* N141I mutation. This also indicates that while there are differences based on genotype in basal conditions, these differences are subtle and may demonstrate more substantial differences in response to external stimulation, like we saw with WNT3A..

### Endothelial Cells with or without PSEN2 N141I Mutation show similar gene expression after Interferon-β stimulation

In order to determine whether the *PSEN2* N141I mutation predisposes to distinct stress responses in ECs, we treated the cells with 50 ng/ml of Interferon-β for 6 hours before harvesting them for long-read RNA sequencing, as well as harvesting untreated controls. Comparing the stimulated cells to their untreated control counterparts, we generated differential expression analysis of these samples, and the volcano plots in **Figures 4A** and **4B** show the genes most upregulated in the IFN-treated samples. To account for variability in cell differentiation, we analyzed the differential expression in the individual replicates, which is shown as a heat map in **4C** and **4D**. Here we see the upregulation of many established IFN-stimulated genes such as *ISG15*, *MK1*, *IFIT1* and *IFIT2*. Responses to IFN-stimulation were similar in REV and SNV iPSC-ECs. In order to determine whether expression differences may lie in certain gene sets as opposed to specific genes, we performed gene set enrichment analysis (GSEA) on the IFN-stimulated samples and compared them to their untreated controls (**Figures 4E** and **4F**). We see that both the control REV cells and the *PSEN2* N141I SNV cells’ most significantly upregulated gene sets were related to viral response, viral defense, and type I interferon stimulation. We did not identify any notable gene sets related to WNT response that differed based on genotype in the IFN-stimulated cells. **Figure 4G** shows a Venn diagram displaying the genes differentially expressed in IFN-stimulated REV and SNV cells, as well as the upregulated cells that are shared between the two. There are 1203 genes shared between the two genotypes after stimulation, with the SNV and REV stimulated cells also exhibiting 605 and 676 uniquely upregulated genes, respectively. After further investigation, we also did not identify WNT-related genes amongst the genotype-specific upregulated genes, indicating that the differences we saw in the earlier figures were not due to gene-level transcriptional changes.

**Figure 4.**
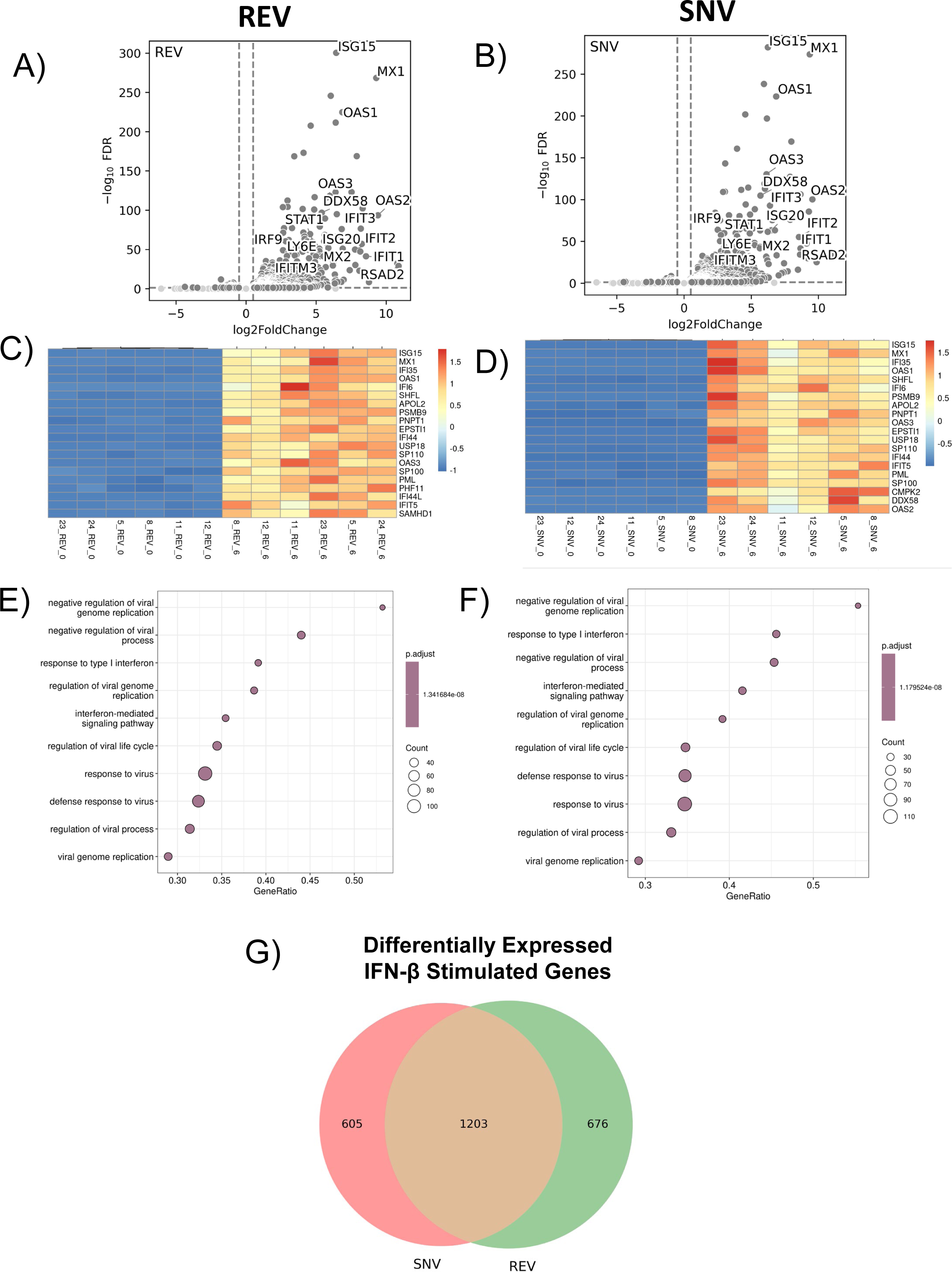
Bioinformatic analysis of IFNβ-stimulated iPSC-derived ECs with (SNV) or without (REV) the *PSEN2* N141I mutation. A)_ Volcano plots show upregulated genes in IFN-stimulated REV and B) SNV iPSC-derived ECs. C) Heat maps comparing differentially expressed genes in REV ECs and D) SNV ECs after IFNβ stimulation. E) Gene Set Enrichment Analysis after IFN stimulation in REV and F) SNV iPSC-derived ECs. G) Venn diagram displaying all genes that are differentially expressed after IFN stimulation in REV and SNV iPSC-ECs. Genes were considered differentially expressed if the |log2(fold change)| > 1.00 and *p<*0.05

### Endothelial cells with PSEN2 N141I mutation show notable upregulation of transposable element expression after IFN stimulation

We next investigated the expression of transposable elements (TEs) in the long-read RNA sequencing data we generated. **Figures 5A** and **5B** show volcano plots of the differentially expressed TEs in IFN-stimulated REV and SNV ECs compared to their untreated control. Some such as *MER5B*, *L1ME1,* and *L1MC5* are uniquely upregulated in the stimulated SNV cells. **Figure 5C** shows a Venn diagram of the TEs upregulated during IFN-stimulation. While 69 TEs are common between the two, the diagram demonstrates how the SNV cells have a large set of 114 TEs that are uniquely expressed during IFN stimulation, while REV only has 35. This major difference in TE expression may explain some phenotypic differences that we see based on *PSEN2* genotype, and deserve further investigation.

**Figure 5.**
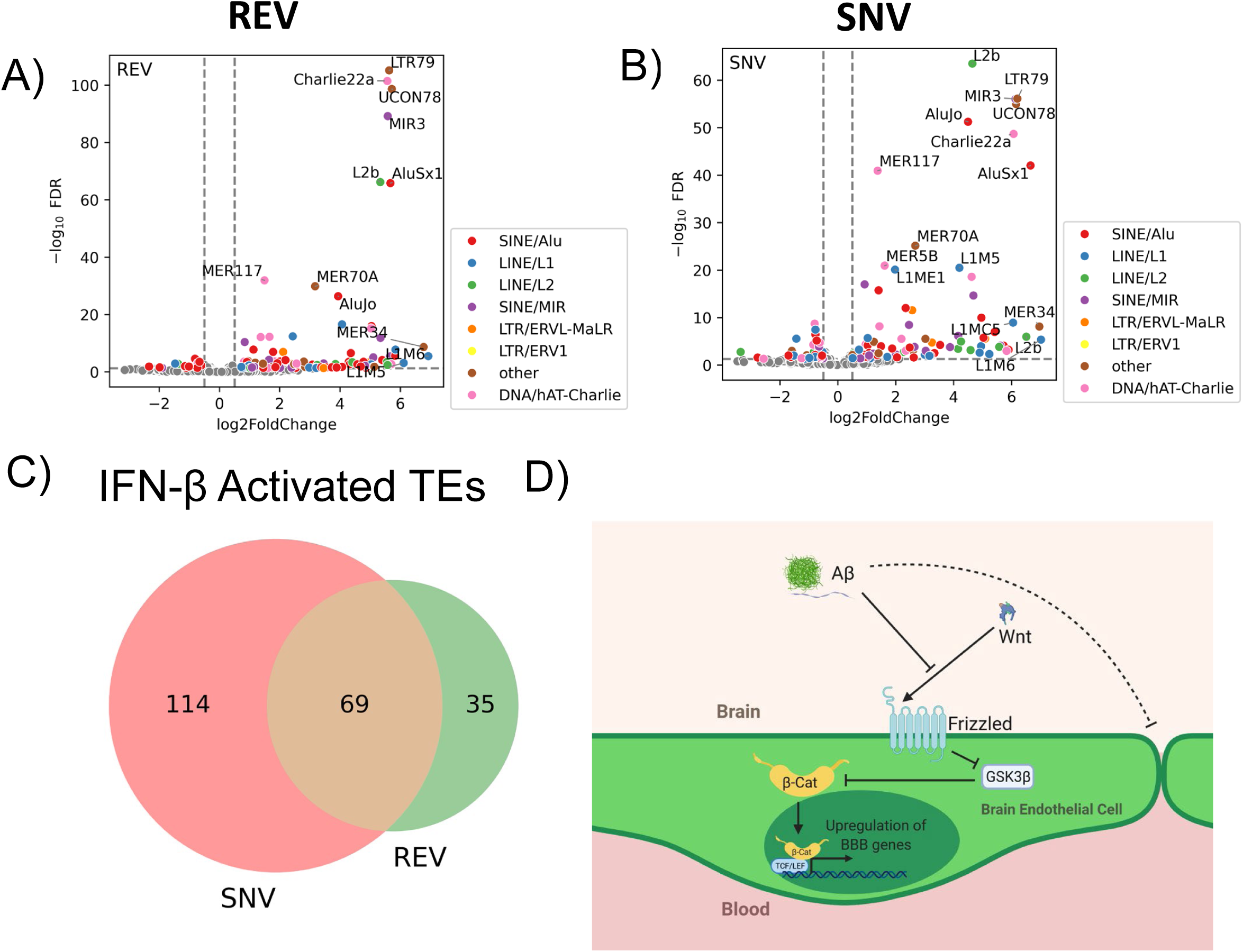
Analysis of transposable element (TE) expression in RNA-sequencing data of IFNβ-stimulated iPSC-derived ECs with (SNV) or without (REV) the *PSEN2* N141I mutation. Volcano plots of TE expression data of IFN-stimulated A) REV and B) SNV iPSC-derived ECs. C) Venn diagram comparing the number shared and unique genes in REV and SNV IFN-stimulated ECs. D) Proposed model of Aβ-induced inhibition of the WNT/β-catenin signalling pathway.

## DISCUSSION

Our data demonstrates a novel model for Aβ-induced BBB disruption through inhibition of the WNT/β-catenin pathways in the brain endothelium. We found that exogenous Aβ exposure led to reduced endogenous expression of β-catenin target genes, inhibition of β-catenin transcriptional activity by WNT, and reduction of the endothelial barrier integrity. Investigating this phenomenon further, we found that Aβ acted within hours of exposure without requiring extensive pre-incubation and demonstrated inhibition of WNT activation also occurred after a 1- hour incubation with the peptides. These phenomena were demonstrated in both the hCMEC/D3 brain endothelial model and iPSC-derived endothelial cells. In iPSC-derived ECs, we found that VE-cadherin protein expression was reduced at cell junctions in cells exposed to Aβ and ECs with genetic backgrounds known to increase endogenous Aβ generation. This supports previous findings of Aβ-induced VE-cadherin inhibition^31^, however, our data suggests that additional junction proteins such as Occludin or Claudin-5 may also respond to Aβ.

Ultimately, the data leads us to the model of Aβ-induced vascular dysfunction shown in **Fig 5D**, where Aβ peptides in the brain parenchyma disrupt WNT stimulation and inhibit β-catenin transcriptional activity leading to reduced BBB properties. We believe this dynamic may represent an early pathological event in the development of Alzheimer’s disease and deserves further investigation as a possible target for AD treatment. Restoring neuronal WNT/β-catenin is also being investigated as a therapeutic target for AD, and the data available indicates this would benefit BBB health as well^20^. Increasing endothelial β-catenin activity could be a therapeutic target for early stages of amyloid generation, hopefully delaying the disease’s progression by maintaining cerebral blood-flow, blood-brain barrier integrity, and endothelial amyloid clearance. Some studies have indicated that targeting brain endothelial β-catenin can restore BBB integrity in mouse subjects^12, 20, 32^. β-catenin activity and barrier protein expression were also reduced in primary rat BECs exposed to amyloid peptides, supporting the model we proposed. Unlike these previous studies, however, we investigated this in the human context using iPSC-ECs, which have great utility for future investigations.

Our study represents an iPSC-based approach to study the effect of Aβ on endothelial β-catenin *in vitro*. Particularly, the use of iPSC-derived endothelial cells provides an opportunity to investigate disease-associated genotypes, such as familial Alzheimer’s disease mutations, and how they influence β-catenin signaling and barrier function *in vitro*. Many models, such as cerebral organoids, have already been utilized with fAD mutation iPSCs to model Aβ deposition and its effect on neuronal function^33, 34^, but the effect of such mutations on the endothelium are far less prevalent in the literature. This currently stands as a blind spot for stem cell based investigation into fAD mutations. Our RNA sequencing analysis did not uncover much in terms of gene expression differences, but the notable upregulation of TE transcripts in the IFN-stimulated *PSEN2* N141I cells may hint at novel mechanisms for how fAD genotypes can influence viral response in the brain, consistent with our previous work^35^.

Our proposed model of Aβ-induced inhibition of the WNT/β-catenin pathway provides a potential mechanism by which the BBB may become dysfunctional in the development of AD. Though previously demonstrated in neurons, our data indicates that restoration of β-catenin as a pharmaceutical target may benefit endothelial function as well as neuronal. Our data coincides with the publication of several new findings that demonstrate that restoring β-catenin can rescue BBB integrity, restore synaptic function, and inhibit generation of Aβ plaques^12, 20^. Previous attempts to generate therapeutic interventions for AD have focused directly on Aβ generation, primarily through amyloid-specific antibodies or by inhibiting beta-secretase to reduce post-translational processing of APP into amyloidogenic Aβ^36^. However, these advancements have struggled to prove efficacious, and thus an alternative target may prove more valuable than investing in drugs that all target the same mechanisms for little benefit.

By generating therapies to target and restore endothelial β-catenin, such a treatment could theoretically restore the BBB’s integrity and ability to clear amyloid from the brain. An engineered WNT ligand that specifically targets Frizzled-4 receptors has been found to restore BBB function and integrity in mouse models of ischemic stroke^32^, indicating there is promise for such an intervention. The key follow-up to support this notion would be to utilize that engineered ligand in an Aβ-generating AD mouse model to determine whether such an intervention would restore the BBB and reduce amyloid plaque load.

Future experiments investigating the effect of fAD mutations on the brain endothelium could provide new insights into the dynamics of AD development. In our hands, we found subtle, though significant changes in the phenotype of basal *PSEN2* N141I compared to their controls. Experiments that stimulate the cells with either WNT, to follow up the previous figures, or IFN-β may demonstrate greater differences than basal. We saw both endogenous β-catenin and WNT-activated catenin was inhibited by Aβ and thus such a dynamic would be worth considering in our model. We could also use our vascularized organoid model with fAD lines and controls in order to study endothelial dysfunction specially in the context of neuronal Aβ exposure. As the major producers of the peptide, the *PSEN2* N141I mutation in neurons may better emulate the context of the BBB in an fAD patient brain, compared to the modest Aβ generation we saw in ECs.

## METHODS

### iPSC Culturing

CRISPR-modified iPSC cell lines with or without the PSEN N141I mutation were obtained as frozen vials from Jackson Laboratory. The SNV/Wildtype line (The Jackson Laboratory #JIPSC1052) and its Revertant/Wildtype control (JIPSC1054) were both cultured in 6-well plates coated with Synthemax II-SC Substrate (Corning #3535) and fed with StemFlex Medium (ThermoFisher Scientific #A3349401). All cell lines were split onto new plates for continued culturing with ReleSR (STEMCELL Technologies #100-0483) at a 1:6 ratio.

The 273 iPSC cell lines were obtained from the Ong Lab at University of Illinois at Chicago. Cells were grown on 6-well cell culture plates coated with Matrigel (Corning #354277). Cells were fed daily with mTeSR1 media (StemCell Technologies #85857). When cells were grown to approximately %70 confluency, they were split for further culturing onto another Matrigel-coated plate using ReLeSR (StemCell Technologies, #100-0483) at a 1:6 ratio.

### hCMEC/D3 cell culturing

The brain microvascular endothelial cell line hCMEC/D3 was obtained from Millipore-Sigma(#SCC066) as frozen vials. The cells were cultured in 75cm^2^ flasks coated with Rat Tail Collagen Type I (Millipore-Sigma #08-115) and fed with EndoGRO-MV Complete Culture Media (Millipore-Sigma #SCME004) supplemented with recombinant human bFGF (R&D #233-FB-010/CF) to a final concentration of 1ng/mL. Cells were split for continued culturing or experimentation using 0.05% Trypsin-EDTA (ThermoFisher Scientific #25300054).

### iPSC differentiation into CD144+ endothelial cells

iPSCs were differentiated into endothelial cells using a previously described protocol^28^. The cells were split using Accutase (STEMCELL technologies, #07920) to break colonies into single cells for more efficient differentiation onto culture plates coated with Geltrex (ThermoFisher Scientific, #A1413202.) The cells were also initially plated with 10µM ROCK inhibitor Y-27632 (STEMCELL technologies, #72303) which was removed on subsequent media changes. The differentiation then proceeded as previously written, following the arterial induction steps described. When the cells were mature and nearing 90% confluency in the plate, cells were again split with Accutase and MACS sorted using anti-CD144 microbeads (Miltenyi Biotec, #130-097-857) onto another Geltrex-coated plate. After nearing 90% confluency in the new plate, the cells were split and sorted with anti- CD144 microbeads and plated onto the final geltrex-coated plates used for the final experiments. These passages and sorting were performed in order to allow the endothelial cells to continue maturing before being used for experiments, around 8-12 additional days of culturing.

### Flow Cytometry

In order to assess the differentiation efficacy of a previously published endothelial cell differentiation protocol^28^, flow cytometry was performed on mature differentiated cells. The cells were collected using Accutase (STEMCELL technologies, #07920), and the cell pellets were washed three times with PBS. After washing, the cells were suspended in PBS containing 0.5% bovine serum albumin and a 1:10 dilution of either anti-CD31 PE conjugated flow antibody (R&D Systems #FAB3567P) or a PE-conjugated antibody isotype control (BD Pharmingen #555749). The cells were incubated with the antibodies for 30minutes on ice.

Afterwards, the cells were washed three times with PBS and centrifugation. Afterwards, the cells were resuspended in PBS with 10%BSA for storage. The cells were fixed with fixation buffer (BioLegend #420801) and flow sorting was performed on a CytoFLEX S Flow Cytometer and analyzed with Kaluza Analysis software (ver 1.5a).

### WNT3A and Amyloid-β treatment of cultured cells

Recombinant human WNT3A protein (R&D 5036-WN-010/CF) was resuspended in 0.1% bovine serum albumin in sterile PBS and added to treatment cell media at a final concentration of 200ng/mL. Amyloid-β 1-42 peptide (rPeptide A- 1166-2) and Scrambled Amyloid-β Peptide (A-1004-2) were suspended in 1% ammonium hydroxide and sonicated to dissolve the peptide per manufacturer’s instructions.

### RNA Isolation and cDNA Generation for RT-qPCR

Cells were collected in TRIzol reagent (ThermoFisher Scientific #15596026) for RNA isolation. RNA was extracted from TRIzol- suspended samples using Direct-zol RNA microprep (Zymo Research #R2061). Isolated RNA was eluted in water and used to generate cDNA using the High-Capacity cDNA Reverse Transcription Kit (Applied Biosystems #4368813). Quantitative PCR was performed in 10µl reactions using SYBR Green PCR Master Mix (Applies Biosystems #4309155) using the manufacturer’s recommended thermocycling protocol.

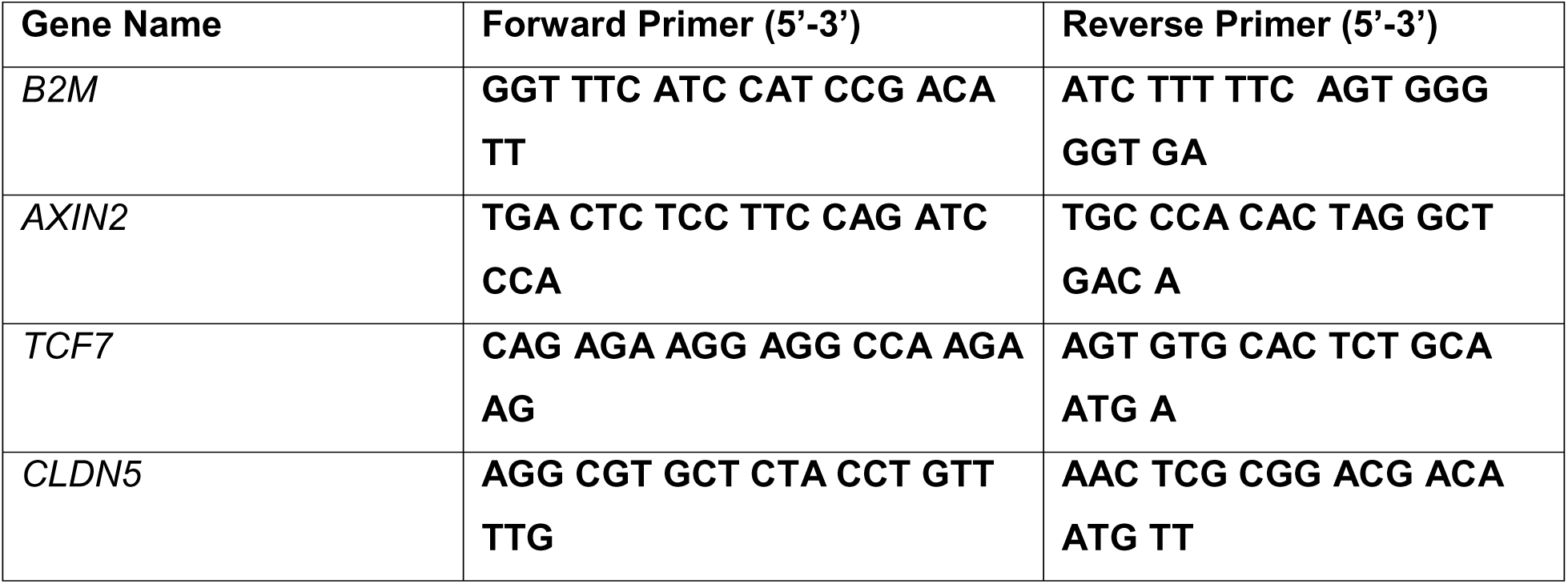

### β-Catenin Activity Luciferase Reporter Assay

hCMEC/D3 brain endothelial cells were grown to confluency in 6-well culture plates and transfected with 1µg of a β-Catenin luciferase reporter plasmid (M50 Super 8x TOPFlash - Addgene #12456) and 35ng of Renila reporter plasmid (pRL-TK Promega #E2231) per well. Treatment of cells began 48 hours after transfection, beginning with 1µM Aβ1-42 or scrambled peptide control. 6 hours before the cell lysate was collected, the cell media was refreshed with either WNT3A (200ng/mL) or vehicle control. 100µl of cell lysate from each sample were used with the Dual-Luciferase Reporter Assay System (Promega #E1910) to measure relative β-catenin activity.

### Immunofluorescence

Cells used for immunofluorescent staining were first plated onto glass coverslips coated with either Rat Tail Collagen Type I (Millipore-Sigma #08-115) for hCMEC/D3 cells or Geltrex matrix (ThermoFisher Scientific #A1413201). Cells were fixed in 4% paraformaldehyde for 10 minutes and permeabilized with 0.25% Triton-X in PBS for 10 minutes prior to antibody incubation. Fixed cells were blocked for 1 hour with 2% BSA solution in PBS. Primary antibody incubation occurred overnight at 4°C with 1:500 647-conjugated anti-VECAD (BD Pharmingen #561567). Cells were mounted on glass slides with VECTASHIELD Antifade Mounting Medium (Vector Laboratories #H-1800). Stained cells were imaged on Zeiss LSM 710 Confocal Laser Scanning Microscope. Each sample had 6-8 images taken across the coverslip, each containing at least 3 full cells in the field of view. Mean fluorescence intensity was measured in ImageJ FIJI using automatic Otsu thresholding to generate a mask for specifically measuring the fluorescence intensity at cell junctions. Cell areas were measured by manually tracing the cell barriers in ImageJ FIJI.

### Interferon-β stimulation and Long-Read RNA Sequencing

iPSCs with (“SNV”) or without (“REV”) the *PSEN2* N141I mutation were differentiated as previously described. Mature cells were then treated with 50 ng/mL Interferon-β (Peprotech #300-02) for 6 hours. Treated and untreated control cells were collected with TRIzol reagent and purified using Direct-zol RNA microprep (Zymo Research #R2061). Sequencing libraries were generated from mRNA using a Oxford Nanopore cDNA-PCR v14 multiplex kit with the standard protocol. Libraries were sequenced on Oxford Nanopore Promethion R10.4 flow cells targeting 20 million reads per sample. Basecalling was done with Guppy using model dna_r10.4.1_e8.2_400bps. Reads were aligned to the telomere-to-telomere genome (GCA_009914755.4) using Minimap2^37^ with flags - ax splice. The transcriptome was annotated with stringtie2 using the GCA_009914755.4 annotation as the reference. The fastq read data was realigned against the transcriptome using Minimap2 with flags -N 100 -ax map-ont. Counts were generated using salmon quant^38^ with flags -l A –ont. Deseq2^39^ was used for differential expression analysis.

### Transposable Element Analysis

GCA_009914755.4 was annotated using RepeatMasker^40^ using the complete Dfam 3.8 database^41^ and RepBase^42^ 20181026 edition with HMMER^43^ used as the search model. All repeats were included in the initial search and then filtered to contain only transposable elements. The TE annotations were combined with the complete GCA_009914755.4 annotation prior to use for transcript counting in order to account for false positives due to gene-TE annotation overlap. FeatureCounts^44^ was used to generate TE counts from the genome alignments and counts from overlapping annotations were discarded.

### Statistical Analysis

All data here are represented as means+/-SEM of at least 3 independent replicated experiments. Shapiro-Wilk test was performed to determine normal distribution for parametric analysis. Normally distributed data was then analyzed with student’s *t* test, repeated measures one-way ANOVA, or repeated measures two-way ANOVA. For repeated measures ANOVA comparison, post-hoc comparisons were made with Bonferroni’s multiple comparisons test and sphericity is assumed. Results of individual multiple comparisons are reported in the figure legends. All statistical analysis was performed using Prism 10, by Graphpad Software.

Significance values are shown in figures as follows: *\*p*<0.05, *\*\*p*<0.01, *\*\*\*p*<0.001, *\*\*\*\*p*<0.0001, and *ns p>0.05*.

## Additional Information

The authors have declared that no conflict of interest exists.

## Acknowledgments

The studies were supported by NIH grants R01-HL163978(J.R.), R01-HL152515 (to JR), T32- HL139439 (to M.A.S.), F31-AG090005 (to M.A.S), T32-HL007829 (to J.W.L),.

