## Supplementary figures and images for "Inhibition of β-catenin signaling by Amyloid-β in endothelial cells impairs vascular barrier integrity"

### Supplementary Figure 1

**Supplemental Figure 1**

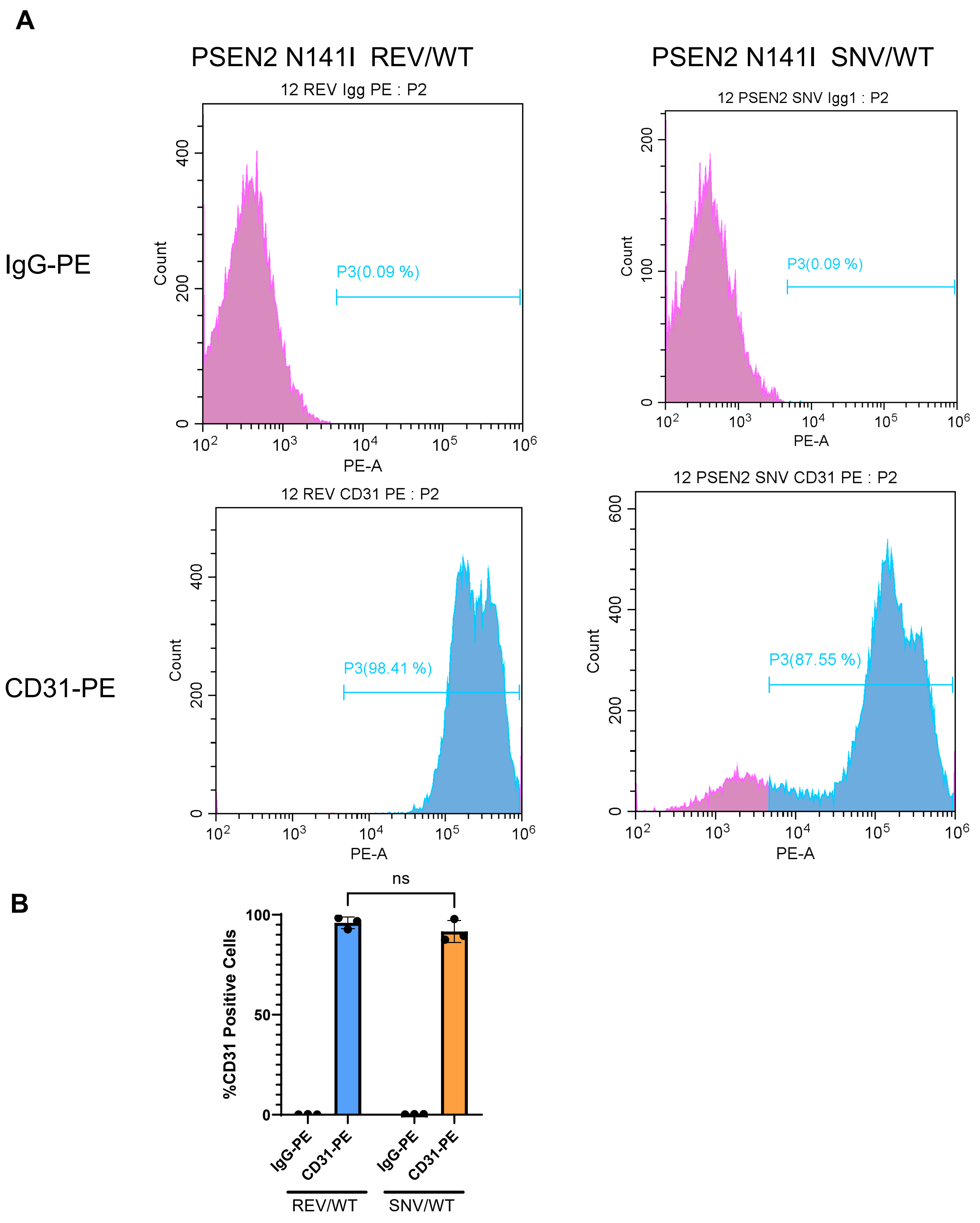
